# Exogenous disturbances and endogenous self-organized processes are not mutually exclusive drivers of spatial patterns in macroalgal assemblages

**DOI:** 10.1101/2020.08.25.265967

**Authors:** J. He, L. Rindi, C. Mintrone, L. Benedetti-Cecchi

**Affiliations:** Department of Biology, University of Pisa, CoNISMa, Via Derna 1, Pisa, Italy; Marine Science and Technical College, Zhejiang Ocean University, Zhoushan City, Zhejiang, China

**Keywords:** spatial patterns, self-organization, scale-invariance, local-scale interactions, power-law distribution, rocky intertidal systems

## Abstract

Complex spatial patterns are common in coastal marine systems, but mechanisms underlying their formation are disputed. Most empirical work has focused on exogeneous spatially structured disturbances as the leading cause of pattern formation in species assemblages. However, theoretical and observational studies suggest that complex spatial patterns, such as power laws in gap-size distribution, may result from endogenous self-organized processes involving local-scale interactions. The lack of studies simultaneously assessing the influence of spatially variable disturbances and local-scale interactions has fuelled the idea that exogenous and endogenous processes are mutually exclusive explanations of spatial patterns in marine ecosystems. To assess the relative contribution of endogenous and exogenous processes in the emergence of spatial patterns, an intertidal assemblage of algae and invertebrates was exposed for 2 years to various combinations of intensity and spatial patterns of disturbance. Localized disturbances impinging at the margins of previously disturbed clearings and homogenous disturbances without any spatial pattern generated heterogeneous distributions of disturbed gaps and macroalgal patches, characterized by a truncated or a pure power-law scaling. Spatially varying disturbances produced a spatial gradient in the distribution of algal patches and, to a lesser extent, also a power-scaling in both patch- and gap-size distributions. These results suggest that exogenous and endogenous processes are not mutually exclusive forces that can lead to the formation of similar spatial patterns in species assemblages.

## Introduction

Complex spatial patterns, such as power laws and irregular distribution of patches are pervasive features of marine ecosystems ranging from mussel beds to seagrasses meadows and coral reefs (Wootton 2001, van der Heide et al. 2010, Edwards et al. 2017). Spatial patterns may have important implications for ecosystem functioning in terms of increased resistance to extreme events and the maintenance of biodiversity (De Paoli et al., 2017; Liu et al., 2014). Although a considerable amount of research has been devoted to understand complex spatial patterns, the processes underlying their formation and persistence are still poorly understood (Rietkerk and van de Koppel, 2008; Tamburello et al., 2013).

Natural disturbances (i.e. discrete events such as storms that remove biomass from a system) are major forces generating spatial and temporal heterogeneity in marine ecosystems (Sousa 1979, Paine and Levin 1981, Dayton et al. 1992). Disturbances can be categorized according to different traits, including frequency, intensity, duration, spatial extent and return time, which combine in various ways to define a wide range of disturbance regimes (Pickett and White 1985). Experimental studies have shown various effects of changing the temporal regime of disturbance on populations and assemblages, either by manipulating the frequency of events or their temporal clustering, sometimes in combination with changes in mean intensity (Johnston and Keough 2002, Bertocci et al. 2005, Benedetti-Cecchi et al. 2006, García Molinos and Donohue 2011). Most of these studies have focused on temporal patterns of disturbance, whereas spatial disturbances have received much less attention. This is surprising, since spatial heterogeneity is pervasive in ecological systems and can be observed both in physical and biological variables. For example, Denny and colleagues (2004) have shown how temperature, wave action and the abundance of sessile organisms followed a ‘red-noise’ spatial pattern with variance increasing continuously in space on a rocky intertidal system in California. Motivated by these observations, Tamburello et. al (2013) exposed algal canopies to spatially autocorrelated, ‘red-noise’ disturbances showing how the spatial characteristics of the perturbations influenced the distribution of the turf-forming algae that developed in response to canopy loss. Despite these experimental insights, our understanding of the extent to which spatially variable disturbances translate into spatial patterns in species assemblages is still limited (Denny et al. 2004, Tamburello et al. 2014).

The theory of self-organization has offered ecologists an alternative paradigm to explain the emergence of spatial patterns in ecosystems in the absence of a spatially-structured disturbance operating at large scales (e.g., a spatial gradient) (Pascual and Guichard 2005, Solé and Bascompte 2012). This theory proposes that complex spatial patterns may emerge as a result of the interactions between small-scale antagonist processes, such as predator-prey and host-parasitoid interactions or disturbance-recovery dynamics (Solé and Bascompte 2012). A well-known example of a self-organized system is provided by rocky intertidal mussel beds, where wave action may create gaps that recover through the lateral encroachment of adult mussels from the margins (Guichard et al. 2003). Edges of newly formed gaps are unstable and more susceptible to be furtherly disturbed than other areas in the bed (Denny 1987). These small-scale interactions, between recovery and disturbance, may propagate trough the mussel bed, producing a characteristic scale-invariant pattern in the gap-size distribution (Guichard et al., 2003).

The interplay between small-scale positive interactions and disturbance may also scale-up to produce complex spatial patterns in marine ecosystems. For instance, on the Pacific coast of North America, individuals of *Mytilus californianus* form a complex network of positive interactions with neighbouring conspecifics and other sessile organisms, which self-organizes into a large-scale spatial pattern in gap-size distribution (Wootton 2001). Spatial patterns in self-organized ecosystems reflect emergent properties that cannot be predicted from the information contained in the external drivers (Heylighen 2002). The emergent feature and qualitative flag of systems governed by small-scale interactions is the scale invariance in patch-size distributions, which means that there is no characteristic patch (or gap) size in the system (Pascual et al. 2002, Kéfi et al. 2011). This scale-invariant pattern can be well described by a power-law or by alternative heavy-tailed functions, such as truncated power-law or exponential functions (Meloni et al. 2017).

Despite the existence of a vast literature on spatial patterns, studies simultaneously investigating the contribution of spatially structured disturbances and self-organization in the formation of spatial patterns remain rare. There are at least two reasons behind this dichotomy. First, theoretical studies on spatial self-organization often assume that disturbance operates homogenously in space, without any particular spatial structure (Koppel et al. 2005, Kéfi et al. 2007, Meloni et al. 2017). Second, very few studies have manipulated small-scale interactions to assess whether these processes scale-up to generate complex spatial patterns. To our knowledge, the only field experiment relating a local process to the formation of a complex spatial pattern is the study of Weerman et al. (2011) on top-down control by herbivores on spatial pattern formation in mudflat biofilm. The relative contribution of self-organization and spatially structured disturbances to the formation of complex spatial patterns remains largely unknown.

In this study, we experimentally assessed the relative contribution of exogenous and endogenous processes to the formation of spatial patterns in rocky intertidal assemblages dominated by the brown alga *C. compressa* in the NW Mediterranean. Exogenous processes were imposed to the system as spatially extended disturbances, whereas endogenous processes were imposed as localized perturbations triggering small-scale dynamics of disturbance and recovery at the margins of already disturbed gaps. In the study system, the cover of *C. compressa* can vary due to the presence of gaps produced by intense storms or as a consequence of desiccation stress resulting from prolonged aerial exposure, which occurs during persistent periods of calm sea and high barometric pressure. Most gaps range in size between 10s to 100s of cm^2^, but larger gaps can be observed occasionally after intense storms (Benedetti-Cecchi and Cinelli 1994). Disturbed gaps become colonized by algal turfs or they may be recolonized by *C. compressa*, but the outcomes of disturbance and recovery are spatially and temporally variable (Benedetti-Cecchi and Cinelli 1994, Benedetti-Cecchi et al. 1996).

Preliminary observations indicated that areas of substratum fully covered by *C. compressa* alternated with others in which the alga was sparse and mixed with patches of algal turfs. Several alternative, not mutually exclusive explanations can be proposed to explain these spatial patterns. First, wave action can be the primary force affecting the distribution and cover of *C. compressa* and algal turfs. However, orientation of the coast, aspect of the substratum and possibly other factors can modify the force with which wave action impinge on rocky shore organisms in different places. Therefore, spatial gradients in the structure of assemblages may reflect spatial gradients in the exogenous forcing effects of wave action.

Spatial patterns may also be the result of the interplay of biotic interactions and disturbance. For example, mussels can increase the resistance of *C. compressa* to wave action by providing additional substratum to which the primary fronds of the alga can attach (Benedetti-Cecchi and Cinelli 1995). Therefore, spatial variation in the distribution of mussels may affect the susceptibility of the alga to be dislodged by waves.

Finally, spatial patterns in *C. compressa* stands may also emerge as a result of local-scale processes, regardless of the spatial characteristics of exogenous disturbances (Pascual and Guichard 2005). Self-organization has been documented in mussel beds where the interactions between small-scale disturbance, acting preferentially along the margins of already existing gaps and recovery scale-up to produce a power-law distribution of gap-sizes (Guichard et al. 2003). *C. compressa* has a very limited dispersal ability due to large buoyant zygotes that settle within 20 cm from the released point (Mangialajo et al. 2012). Limited dispersal (local recovery) coupled with local-disturbance operating at the margin of previously disturbed gaps may produce a near or pure scale-free pattern in stands of *C. compressa* and algal turfs.

To examine the alternative explanations outlined above, we exposed patches of *C. compressa* to experimental perturbations with different spatial characteristics: 1) a gradient-like pattern (GRAD), 2) a homogeneous pattern (HOM) and 3) disturbances that operated locally at the margins of previously disturbed gaps (EDGE). Each of these treatments was crossed with two levels of disturbance intensity according to a two-way factorial design. The study also included unmanipulated controls and naturally fragmented patches for reference. Response variables included the percentage cover and spatial autocorrelation of *C. compressa* and algal turfs and the size of disturbed gaps in each treatment. This experiment allowed us to test the hypotheses originating from the alternative explanations outlined above:

*Hypothesis 1* – If wave action or other exogenous perturbations operate along spatial gradients, *C. compressa*, algal turfs and disturbed gaps should also display spatial gradients in the GRAD treatment and these patterns should differ from those observed in the other treatments, at least for one of the two levels of intensity of disturbance examined. Furthermore, patterns in the GRAD treatment should differ from those observed in unmanipulated controls and should become similar to those occurring in naturally fragmented patches.
*Hypothesis 2* – If exogenous disturbances do not operate along spatial gradients, the spatial distribution of *C. compressa*, algal turfs and disturbed gaps should become randomly distributed in the HOM treatment and spatial patterns for this treatment should become similar to those observed in naturally fragmented transects. Furthermore, patterns in the HOM treatment should differ from those in the controls and in the other treatments, at least for one of the two levels of disturbance examined.
*Hypothesis 3* – If self-organization originating from the interplay between disturbance and small-scale processes (e.g. local recovery or biotic interactions) is the main driver of the formation of spatial patterns, the spatial distribution of algal turfs and disturbed gaps should follow a pure or nearly pure power-law distribution in the EDGE treatment, which should become similar to what observed in naturally fragmented conditions. In addition, patterns in the EDGE treatment should differ from those observed in the controls and in the other treatments, at least for one of the two levels of disturbance examined.
*Hypothesis 4* – The hypotheses outlined above are not mutually exclusive and different combinations are possible. In the event that both exogenous disturbances and spatial self-organization are important processes, we expected signatures of self-organization (e.g. power-law distribution) and spatial gradients to be present in the same treatments. We note that none of the experimental treatments may converge to naturally fragmented patches if both exogenous and endogenous processes are important in the system, because our design did not include combinations of experimental perturbations (GRAD + EDGE or HOM + EDGE).

## Materials and Methods

### Study system

This study was done along the rocky shore of Calafuria, few kilometers south of Livorno, Italy (43°31’N, 10°20’E). At heights on the shore ranging between 0 to −0.3 m below the Mean Low Water Level (MLWL), the brown alga *Cystoseira compressa* (Esper) Gerloff & Nizamuddin forms belts that extend for several meters in the alongshore direction, alternating with areas dominated by algal turfs, clumps of mussels, encrusting algae and apparent bare rock. Storms can generate gaps within belts of *C. compressa* ranging in size from tens to hundreds of cm^2^, although larger patches can be observed occasionally after unusually intense storms. Algal turfs are intricate mats of low-lying, structurally simple algae (Connell et al. 2014). In our study system, algal turfs generally include articulate coralline algae, coarsely branched algae, filamentous and thin-tubular or sheet-like algae (Benedetti-Cecchi et al. 1996, Maggi et al. 2009).

### Experimental design

In November 2017 we permanently marked 21 transects of 180 × 30 cm oriented in the alongshore direction in areas fully covered by *C. compressa* along a stretch of coast of 1 km. To assess the response of *C. compressa* and algal turfs to disturbances with different spatial characteristics, transects were manipulated according to a two-way factorial design (Fig. 1). Six transects were randomly assigned to each of three distinct spatial patterns of disturbance: (1) gradient-like disturbances (GRAD) where the probability of disturbance decreased along the longer axis of the transects, (2) spatially homogenous disturbances (HOM), where the probability of disturbance was homogenous along the transects and (3) localized disturbances (EDGE) where disturbances occurred at the margins of previously disturbed gaps. The six transects allocated to each type of disturbance were randomly assigned in replicates of three to one of two levels of intensity: a low level of intensity (L) where 8% of the surface of a transect was disturbed each time (experimental disturbances were repeated four times during the course of the study, see below) or a high level of intensity (H) where 16% of the transect was disturbed. This experimental layout used 18 of the 21 transects; the remaining three transects were left untouched and served as controls. Control transects were fully covered by *C. compressa* with no apparent disturbed gap at the beginning of the study, allowing us to evaluate the influence of natural disturbances on assemblages during the course of the experiment. Three additional transects were established at the beginning of the study within naturally heterogenous patches of *C. compressa*. These transects served as a reference condition to assess which, if any, of the experimental treatments generated spatial patterns in assemblages that resembled those observed in naturally fragmented conditions (Supplementary material Appendix 1 Fig. A1).

**Figure 1.**
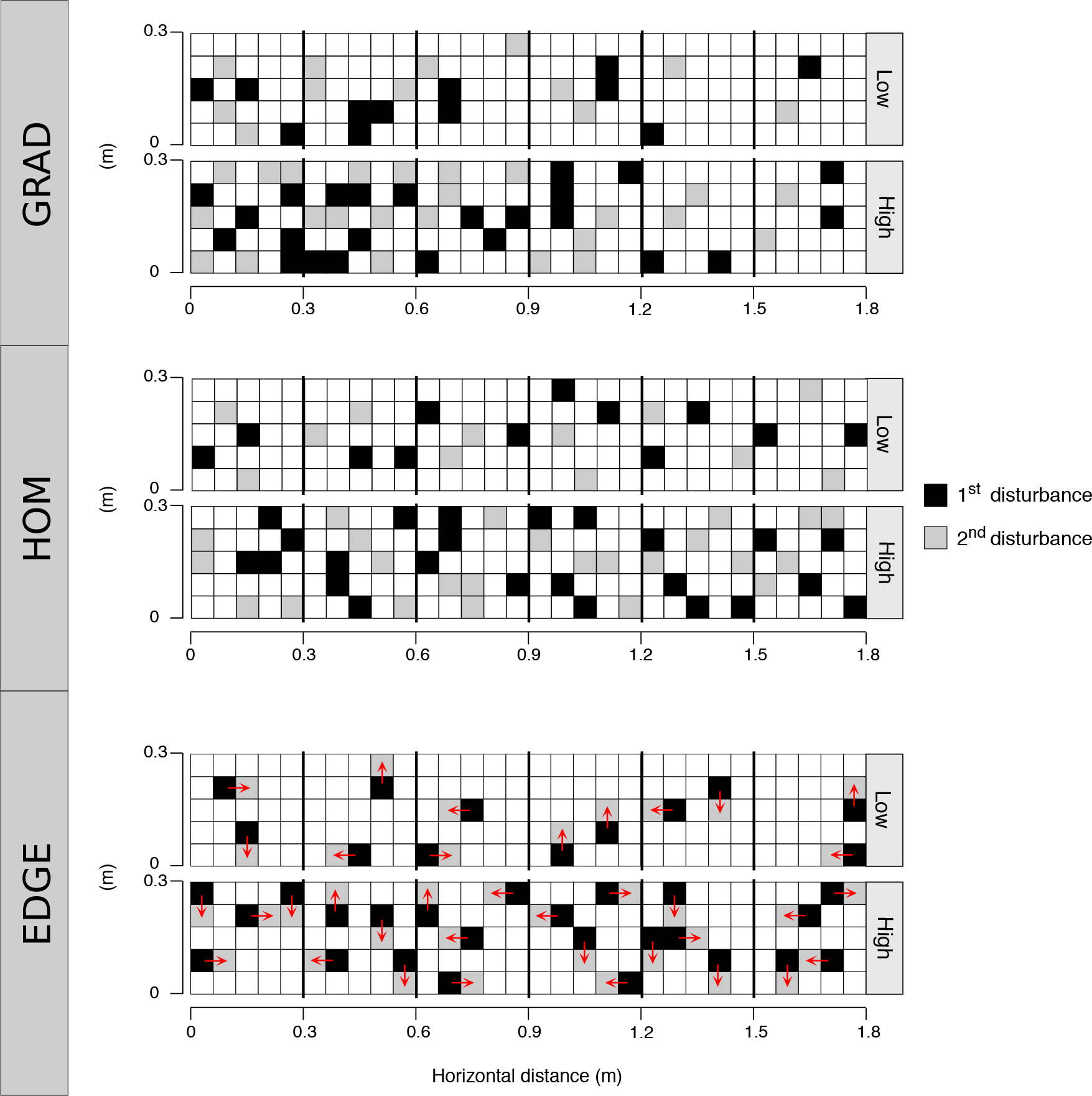
Schematic illustration of experimental treatments. Examples of the three spatial patterns of disturbance (GRAD, HOM and EDGE) are shown for each level of intensity. Experimental transects of six 30 × 30 cm contiguous quadrats were established in areas originally covered by *C. compressa*. The availability of primary space was manipulated to generate three spatial patterns of disturbance: gradient (GRAD), homogenous (HOM), and localized (EDGE). For each spatial pattern, three transects were assigned to each of two levels of intensity: Low and High. Treatments were stratified within fully covered stands of *C. compressa*. Clearings of 6 × 6 cm were distributed along the main axis of each manipulated transect at the start of the experiment. Randomization schemes were used such that clearings were distributed to generate a spatial gradient of disturbance along transects in the GRAD treatment, whereas homogeneously distributed clearings were produced in HOM and EDGE treatments. New randomization schemes were used to distribute clearings at subsequent treatment dates, maintaining a spatial gradient in the GRAD treatment and homogenously distributed disturbances in the HOM treatment. Clearings were distributed homogenously in the EDGE treatment at the first sampling date, but subsequent disturbances were produced at the margins of previously disturbed gaps in subsequent treatments. In the scheme, black and grey squares are gaps produced at the 1^st^ and 2^nd^ disturbance, respectively. Red arrows in the EDGE transects indicate the direction of the expansion of previously disturbed gaps.

Disturbances consisted in the complete removal of erect organisms with hammer and chisel from the substratum in areas of 6 × 6 cm (Supplementary material Appendix 1 Fig. A1c), well within the range of gap sizes generated by natural perturbations (Benedetti-Cecchi and Cinelli 1994). To generate a spatial gradient of disturbance (GRAD treatment) each transect was viewed as a sequence of 6 contiguous quadrats of 30 × 30 cm, each of them receiving a prescribed number of clearings as follows: 6-6-4-4-2-2 clearings for the high intensity treatment (the first and second quadrat in the transect had six clearings, the third and fourth quadrats had four clearings and the fifth and sixth quadrats had two clearings) and 3-3-2-2-1-1 clearings for the low intensity treatment (Fig. 1).

Disturbances were imposed four times during the experiment (Supplementary material Appendix 1 Fig. A2). Clearings were distributed randomly within quadrats in the GRAD treatment and a new randomization scheme was used at each time of disturbance, with the proviso that each quadrat in the transect received the prescribed number of clearings. The same number of clearings used for the GRAD treatment were fully randomized along transects in the HOM and EDGE treatments at the beginning of the experiment. A new randomization scheme was used at each time of disturbance in the HOM treatment, whereas subsequent disturbances were applied at the margins of previously disturbed gaps in the EDGE treatment (Fig. 1). This latter treatment was designed to mimic what observed in mussel beds, where the margins of previously disturbed gaps become susceptible to further disturbances, which is a condition that can lead to the emergence of self-organized patterns in mussel beds (Denny 1987). Therefore, the EDGE treatment was key to quantify the contribution of local-scale disturbance-recovery processes taking place at the margins of disturbed gaps to the emergence of large-scale patterns.

### Sampling and data analysis

All transects were sampled in August 2018 and July 2019, about 4 months after the 2^nd^ and the 4^th^ experimental disturbances and during the peak of vegetative growth of *C. compressa* (Supplementary material Appendix Fig. A2) (Falace et al. 2005). Visual estimates of percentage cover were used to assess the abundance of organisms in each of the 30 × 30 cm plots along the transects. The percentage cover of sessile organisms was estimated with the aid of a 30 × 30 cm plastic frame divided into 25 sub-quadrats of 6 x 6 cm and giving a score from 0 to 4 to each species in each sub-quadrat.

### Correlograms

To describe the spatial patterns that emerged under different disturbance scenarios, we computed the spatial autocorrelation at different distance classes (or *lags*) on the percentage cover of *C. compressa* and algal turfs (Fortin and Dale 2014). We used the 2D anisotropic non-parametric function in R package ‘ncf’ (Bjornstad 2020) to derive correlograms along the major axis of each transect. We derived an average correlogram for each experimental condition. Significance of Moran’s *I* at different spatial *lags* was calculated trough a permutation procedure (*n*=1000). At each iteration, cells of each transects were reshuffled and correlograms were estimated and averaged separately for each treatment. The averaged Moran-*I* at a given lag was deemed as significant when it exceeded the critical values corresponding to the 2.5^th^ and 97.5^th^ percentiles of the null distribution obtained from averaged correlograms within the same treatment. To avoid bias related to the low number of data at large distances, spatial autocorrelation was calculated up to a maximum *lag* of 144 cm (Fortin and Dale 2014).

### Signatures of Self-organization

Power-law patterns are predicted to emerge in self-organized ecosystems, where the interplay between the localized disturbance and recovery scales-up to the whole system generating a scale-invariant pattern (Pascual and Guichard 2005). High-resolution data of percentage cover of *C. compressa* and algal turfs were converted to binary data and analysed separately. We defined as empty those cells (6 x 6 cm) where the cover of *C. compressa* was ≤ 1% and as occupied those cells in which the cover of algal turfs was ≥ 3% (alternative analyses with different binary thresholds for both *C. compressa* and algal turfs yielded qualitatively similar results). Binary data were analysed with the R package ‘raster’ (Hijmans 2019) in order to identify gaps in *C. compressa* stands and patches occupied by algal turfs. A gap in *C. compressa* stands (or a patch of algal turfs) was defined as a set of empty (or occupied) cells sharing at least one side with their neighbours. We described, for each experimental transect, the frequency of gap and patch sizes using an inverse-cumulative distribution, which is the probability that the size of gaps (S) was larger than or equal to a given value *s*, *P*(*S* ≥ *s*), as a function of size (Kéfi et al. 2014, Génin et al. 2018). Note that in contrast to previous studies, this procedure is not based on binning, which ignores the information of gap- (or patch-) size distribution within each bin.

Three alternative functions were fitted to the gap- and patch-size (S) distribution of *C. compressa* and algal turfs: a power-law (*CS*^−λ^), a truncated power-law (*CS*^−λ^ *e*^−*βs*^), where λ is the scaling exponent, *β* is the rate of exponential decay, and C is a constant, and a log-normal 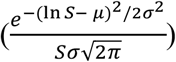 function, where μ and σ are the mean and the standard deviation, respectively (Kéfi et al. 2007, 2011, 2014). The last function represents a “null” model, which is expected when selforganization does not operate and the gap-size distribution is dominated by gaps of a specific size. Transects with less than 4 unique gap (patch) sizes were excluded from the fitting procedure (Génin et al. 2018). Estimates of models parameters were obtained using the Maximum Likelihood estimation method following the Clauset procedure (Clauset et al., 2009). Parameters were then rescaled using a bounded optimization method with the Logit Link function of ‘VGAM’ R package (Yee 2015). The best-fitting model, among those tested, was selected on the basis of the lowest Akaike Information Criterion (AIC) (Clauset et al., 2009). The core steps of the analysis were performed using R package ‘spatialwarnings’ (Génin et al. 2018). Coefficients and confidence intervals are reported in Supplementary material Appendix 1 Tables A1 and A2.

## Results

### Autocorrelation structure

Heat maps of the percentage cover of *C. compressa* showed considerable variation along transects and between sampling dates (Fig. 2, Supplementary material Appendix 1 Figs. A3-A4). Spatial gradients were evident in the GRAD-L and EDGE-H treatments at both sampling dates, whereas *C. compressa* was more patchily distributed in the other treatments. In general, percentage cover declined from August 2018 to July 2019 in all conditions including control and reference transects, the latter showing a more pronounced trend and supporting less cover of *C. compressa* than all the other conditions (Supplementary material Appendix 1 Fig. A5)

**Figure 2.**
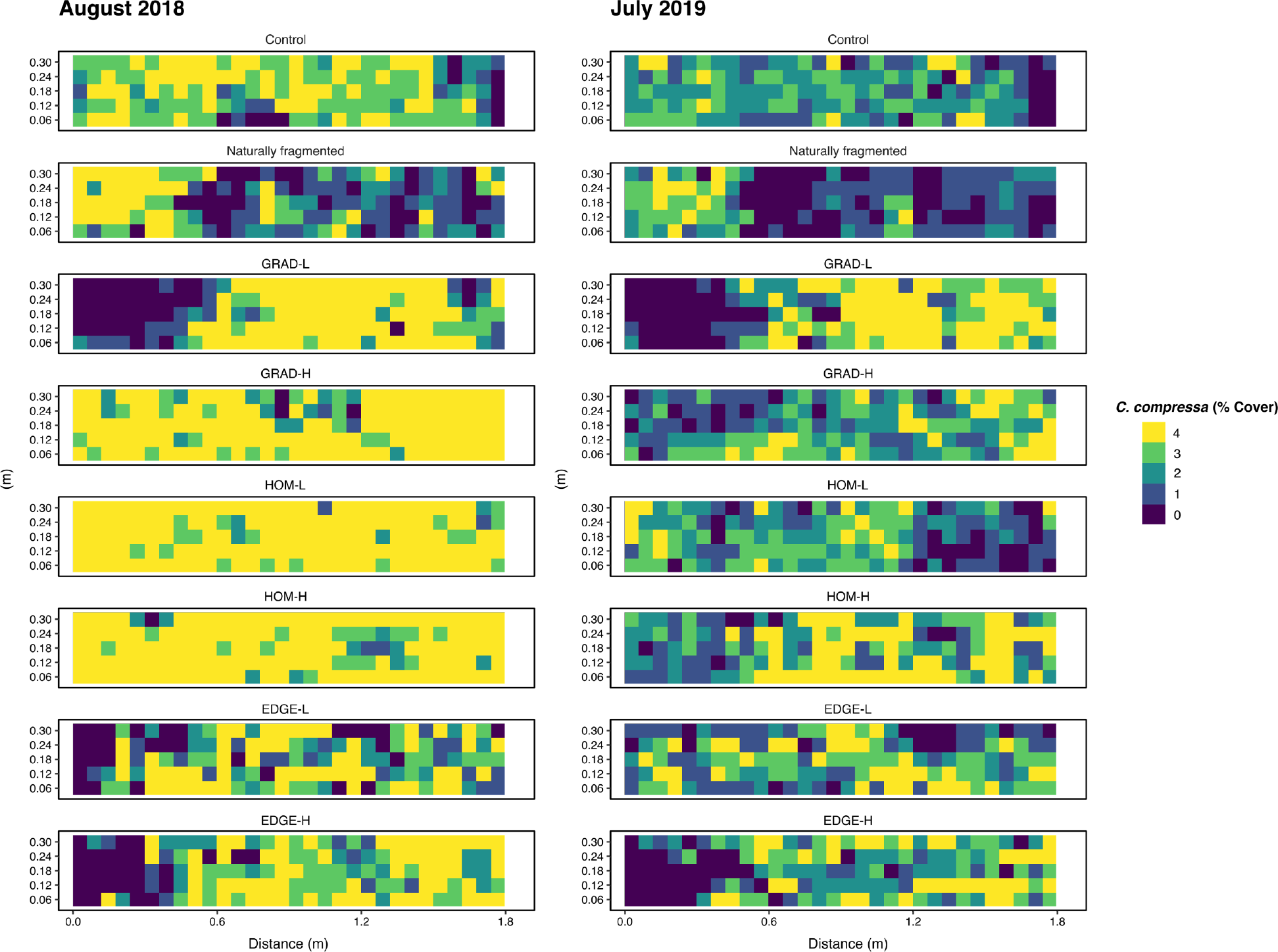
Heat-maps of *C. compressa* cover. Heat-maps of percentage cover data in August 2018 and July 2019 are shown for *C. compressa* using one replicate transect for each experimental condition as an example (see Supplementary material Appendix 1 Fig. A3 for the other transects).

The spatial patterns of *C. compressa* visualized through the heat-maps were supported by correlograms. Both GRAD-L and EDGE-H treatments showed significantly positive correlation at short distances that switched to negative values at larger distances, highlighting strong unidirectional gradients (Fig. 3a,b). These patterns were remarkably similar to those observed in naturally fragmented transects and deviated from those occurring in the controls, which showed a semi-periodic pattern with a transition from positive to negative correlation at short distances and positive correlation raising again at larger distances (Fig. 3a,b). Less strong gradients were observed in the HOM treatment in July 2019 under both levels of intensity of disturbance, but these patterns did not differ substantially from the controls.

**Figure 3.**
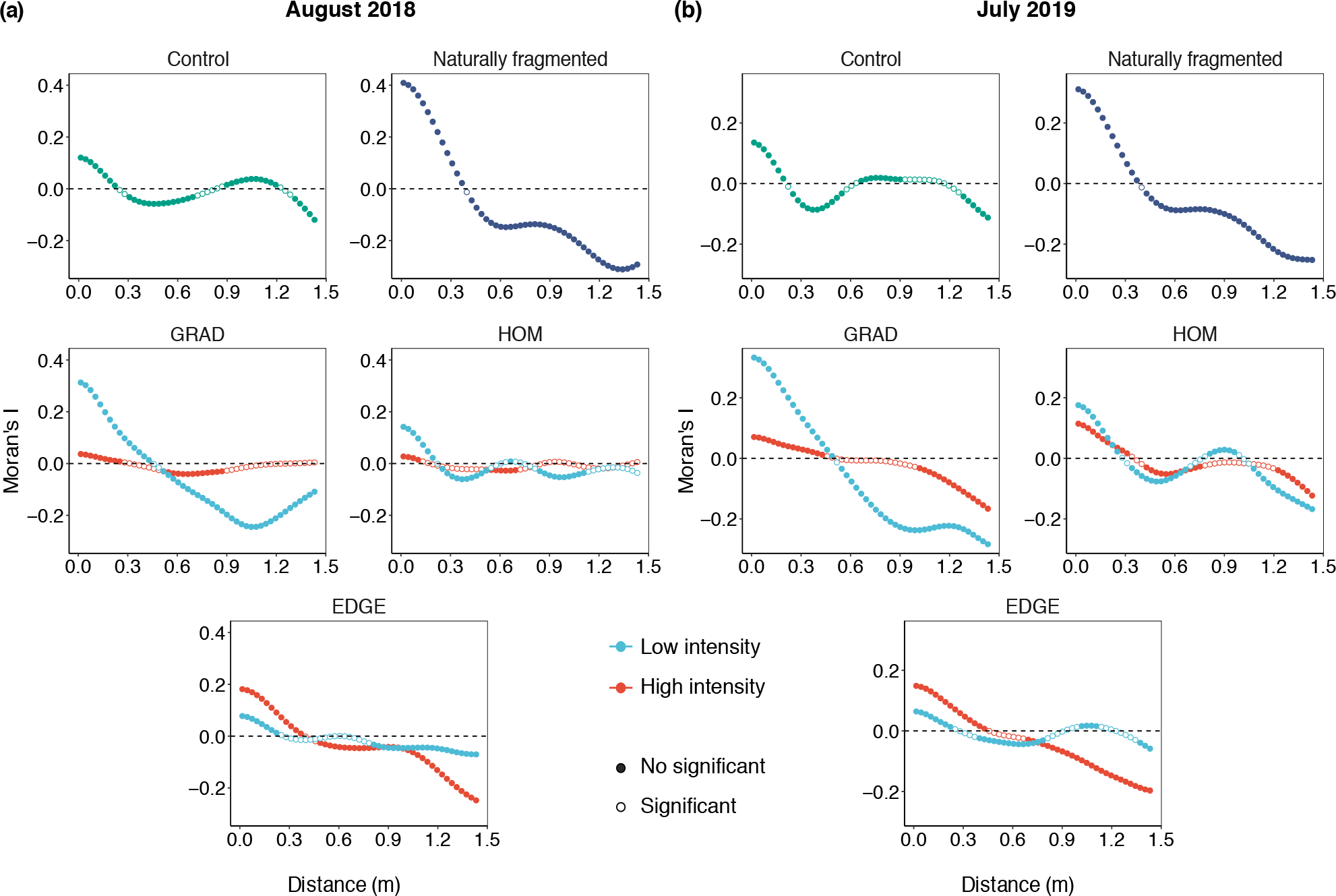
Moran’s *I* correlograms of *C. compressa* cover. Spatial autocorrelation (Moran’s *I*) as a function of distance from one edge of the transect (arbitrarily chosen) in August 2018 (a) and July 2019 (b). Significant values of Moran’s *I* (at α = 0.05) are denoted by filled symbols. Moran’s *I* significance at each distance was tested over 1,000 permutations. Note that in order to improve readability, the y-axis limits are rescaled according to observed values of Moran’s *I*.

The spatial patterns of algal turfs were very similar to those described for *C. compressa* and are reported in Supplementary material Appendix 1 Fig. A6.

### Signatures of self-organization

Pure or truncated power laws in the frequency distribution of patches of algal turfs were observed both in control and in naturally fragmented transects in August 2018 and in July 2019 (Fig. 4, Supplementary material Appendix 1 Table A1). Also, the distribution of gap sizes within *C. compressa* followed a power law in control and in naturally fragmented transects, although the enlargement and coalescence of gaps precluded a formal analysis of fragmented transects in July 2019 (Supplementary material Appendix 1 Fig. A7, Table A2).

**Figure 4.**
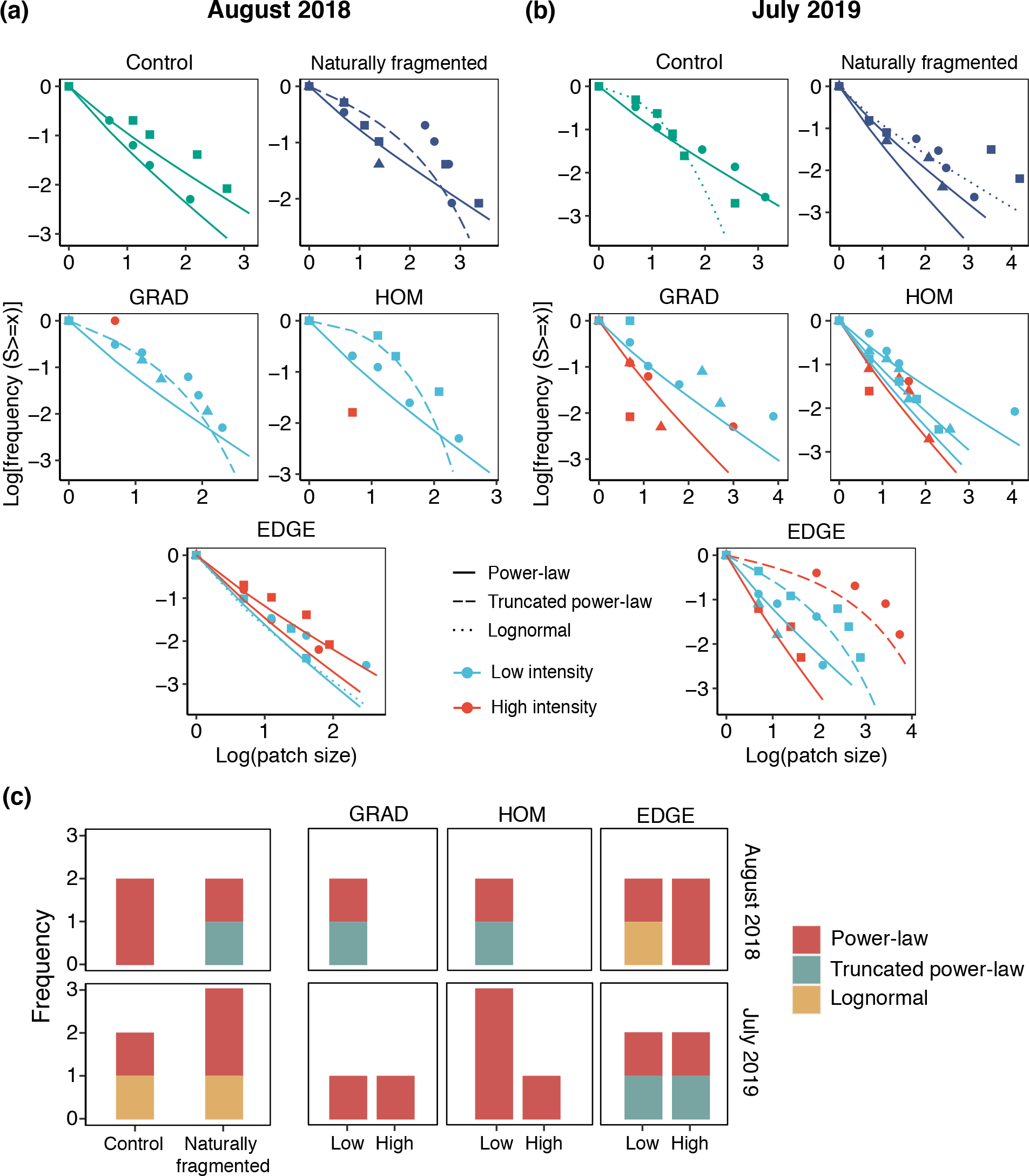
Frequency distributions of algal turfs patch sizes in experimental transects. Plots of log-inverse cumulative distribution of patch sizes - log-transformed frequency of patches larger than a certain size (*s*) - as a function of the log of the patch size in August 2018 (a) and July 2019 (b). Lines denote the three alternative functions: power-law (solid), truncated power-law (dashed) and log-normal (dotted). Symbol shapes denote replicate transects (n=3). Transects with less than four distinct patch sizes were excluded from the fitting procedure. (c) Absolute frequency of AIC-selected functions (among those tested) in different experimental conditions in August 2018 and July 2019. Colors denote the three alternative distributions: power-law (red), truncated power-law (green) and log-normal (yellow).

Pure or truncated power laws were selected more often that the lognormal as model fits of the frequency distribution of patches of algal turfs in experimental transects (Fig. 4a,b, Supplementary material Appendix 1 Table A1). Power laws were visible in transects of the HOM treatment for both levels of intensity of disturbance and this pattern was particularly pronounced for the distribution of gap sizes (Fig. 4, Supplementary material Appendix 1 Fig. A7). Also transects that displayed strong spatial gradients, such as in the GRAD-L and HEDGE-H treatments followed a power law distribution (Fig. 4, Supplementary material Appendix 1 Fig. A7). Due to the limited number of different gaps or patches of unique size, a statistical analysis of gap- and patch-size distributions was not possible for some of the transects exposed to high intensity of disturbance (Fig. 4, Supplementary material Appendix 1 Fig. A7).

## Discussion

We evaluated the importance of exogenous (disturbance) and endogenous (self-organization) processes in the emergence of spatial patterns through a field experiment in a rocky intertidal system dominated by the brown alga *C. compressa*. Spatial gradients in the distribution of *C. compressa* and algal turfs originated in response to unidirectional disturbances of low intensity (GRAD-L), with patterns mimicking those observed in naturally disturbed transects. Spatial gradients also emerged in the treatment designed to reproduce local scale interactions of disturbance and recovery typical of self-organized processes (EDGE-H) (Pascual and Guichard 2005). The frequency distribution of patches of algal turfs and disturbed gaps followed pure or truncated power laws in control, naturally fragmented and experimental transects, including those exhibiting strong spatial gradients (GRAD-L and EDGE-H) and those in which perturbations were distributed randomly in space (HOM). Collectively, these results suggested that the formation of natural spatial pattern was not the outcome of a single dominant process within *C. compressa* stands. Indeed, results provided support to *hypothesis* 4 (see Introduction), showing how spatially-structured disturbances and self-organization may jointly shape spatial patterns in these macroalgal assemblages. Our results have implications for how structuring processes are inferred from observations and model fits. Specifically, power laws may originate in response to directional gradients of disturbance and are not an exclusive signature of self-organization. Conversely, spatial gradients in assemblages may result from spatially random perturbations and from local disturbance-recovery processes that underly self-organization (Guichard et al. 2003, Pascual and Guichard 2005).

The ability of EDGE-H disturbances to generate spatial gradients was intriguing. Perturbations were distributed randomly in space at the start of the experiment and subsequent disturbances were imposed at the edges of existing gaps. A possible explanation for the occurrence of spatial gradients in the EDGE treatment is that perturbations interacted with some physical or biological features of the environment. For example, a previous study showed how the mussel *Mytilus galloprovincialis* could increase the resistance of *C. compressa* to wave action by providing additional substratum for the primary fronds of the alga, thereby reducing whiplash and increasing resistance to dislodgement (Benedetti-Cecchi and Cinelli 1995). Thus, the effect of disturbing the edges of previously disturbed gaps likely differed in areas colonized by *M. galloprovincialis* compared to those where mussels were absent. Since mussels formed clumps in the study area *(personal observation;* Supplementary material Appendix 1 Fig. A1d) their distribution may have modulated the response of macroalgae to experimental disturbance in the EDGE treatment.

The emergence of gaps in control transects indicated that natural disturbances were operating during the experiment and their effects must be considered for a correct interpretation of the results. HOM transects exhibited periodic patterns of spatial autocorrelation analogous to those observed in controls. Thus, natural perturbations operated randomly in space, at least when impinging over fully covered patches of *C. compressa*. However, naturally fragmented transects exhibited strong spatial gradients, indicating that the effects of natural perturbations could change over time depending on the degree of algal loss. Fragmented transects had larger gaps and much less cover of *C. compressa* than controls throughout the study. As the cover of *C. compressa* decreased in response to disturbance, randomly distributed gaps may have coalesced, interacting with other processes to produce more complex spatial structures as discussed for the EDGE treatment.

The similarity of the spatial gradients originating in the GRAD-L treatment to those observed in naturally fragmented transects suggested that spatial heterogeneity in assemblages could also originate as a direct consequence of heterogeneous disturbances. A similar result was obtained in a previous field experiment showing how spatially autocorrelated (red-noise) disturbances producing gaps in the canopy-forming alga *C. amentacea* induced a shift in the distribution of algal turfs that closely tracked the spatial structure of experimental perturbations (Tamburello et al. 2013). Altogether, these results contributed to reinforce the view that exogeneous and endogenous processes are not mutually exclusive explanations for the spatial organization of these macroalgal assemblages.

Intense disturbances did not amplify the spatial patterns observed at lower intensity of disturbance. In general, heavily disturbed transects had a limited number of unique gap sizes and an excess of small gaps that in several instances prevented the fitting of a power law distribution. This was surprising, since large gaps were produced when imposing high levels of disturbance intensity experimentally. For some reasons, *C. compressa* may have been able to colonizae large gaps faster than smaller ones. Similar patterns have been documented in other studies, showing how algal colonization in small gaps may be inhibited by competition, grazing and lack of recruitment, none of which have been explcitly investigated in the present study (Sousa 1984, Farrell 1989). Nevertheless, the cover of *C. compressa* was always larger in experimental than in naturally fragmented transects, indicating that our experimental treatments had not yet reproduced the cumulative levels of disturbances that could occur in nature. As in any manipulative studies, our results were contingent on the duration of the experiment and more time was probably needed for spatial patterns analogous to those originating in naturally fragmented transects, with a sparse cover of *C. compressa*, to emerge under intense experimental disturbance. However, our experiment lasted two years, long enough to examine disturbance-recovery processes in this system (Benedetti-Cecchi and Cinelli 1994, Benedetti-Cecchi et al. 1996).

Pure or truncated power-laws were pervasive in our experiment, emerging in all manipulated and unmanipulated conditions. This suggested that both spatially structured and unstructured disturbances promoted the emergence of power-law patterns in *C. compressa*. For example, a power law distribution may have emerged in the GRAD treatment from the combination of numerous small gaps originating from the weakly disturbed areas of the transect with few large gaps originating from the heavily perturbed region. Conversely, small-scale processes of disturbance and recovery likely promoted the origin of power laws in the HOM and EDGE treatments. Recruitment limitation may be part of the explanation. *C. compressa* has a very limited dispersal ability due to large buoyant zygotes that settle within 20 cm from the released point (Mangialajo et al. 2012). Limited dispersal (local recovery) coupled with local disturbances operating at the margin of previously disturbed gaps, may have been the antagonistic processes behind the scale-free patterns observed in this study. Interactions between local antagonistic processes are responsible for complex spatial patterns in coastal ecosystems, such as mussel beds and mudflats colonized by diatom biofilm (Pascual et al. 2002, Weerman et al. 2012, Saravia et al. 2012). For instance, in mudflats, scale-free patterns in the patch-size distribution of diatom biofilm originate from the interactions between sediment accumulation induced by the growth of biofilm and sediment drainage due to water currents (Weerman et al. 2012). Notably, we cannot rule out that local antagonistic processes may have also operated in GRAD transect amplifying the observed power-law patterns.

Gap- and patch-size distributions have been widely used to infer prevalent processes in ecosystems as well as the proximity to a critical threshold (Rietkerk et al. 2004, Kéfi et al. 2007, Weerman et al. 2012, Scheffer et al. 2015). For instances, in drylands, patch-size distributions characterized by a power-law scaling have been suggested to be the consequence of facilitation mechanisms, whereas a truncated power-law relationship in patch-size distribution has been associated with an excess of stress or competition (Kéfi et al. 2007, 2011). Our results showed how power-law patterns could emerge as a result of disturbances operating along a spatial gradient. Although patch-size indicators may provide a valuable resource in systems not amenable to experimental manipulation, caution must be paid in relying on patch-based indicators to infer dominant processes in ecosystems, because, as shown in this study, a power-law pattern may also emerge in response of spatially-structured disturbances. Furthermore, our results emphasize the importance of considering the intensity of disturbance in modulating the emergence of scale-invariant distributions, in agreement with theoretical studies showing how scale-free patterns in patch-size distribution may occur only within a specific range of disturbance intensity (Pascual et al. 2002).

In conclusion, our study emphasized the need to consider both exogenous and endogenous processes as simultaneously operating forces underpinning the formation and maintenance of spatial patterns in macroalgal assemblages. The transient nature of gap- and patch-size distributions observed both in experimental and in naturally fragmented transects strongly indicated that spatial and power-law patterns represented a dynamic feature rather than a static property of these assemblages. The mechanisms explaining our results, including environmental heterogeneity, biological interactions and disturbance-recovery dynamics are pervasive in nature and we expect our results to apply broadly to terrestrial and aquatic ecosystems. Our findings should thus motivate further research exploring the compound effects of endogenous and exogenous processes, building on the strength of those studies that have elucidated the contribution of disturbance and self-organization as key forces shaping spatial pattern in ecosystems (Guichard et al. 2003, Pascual and Guichard 2005, Kéfi et al. 2011).

