## Supplementary information for "Exogenous disturbances and endogenous self-organized processes are not mutually exclusive drivers of spatial patterns in macroalgal assemblages"

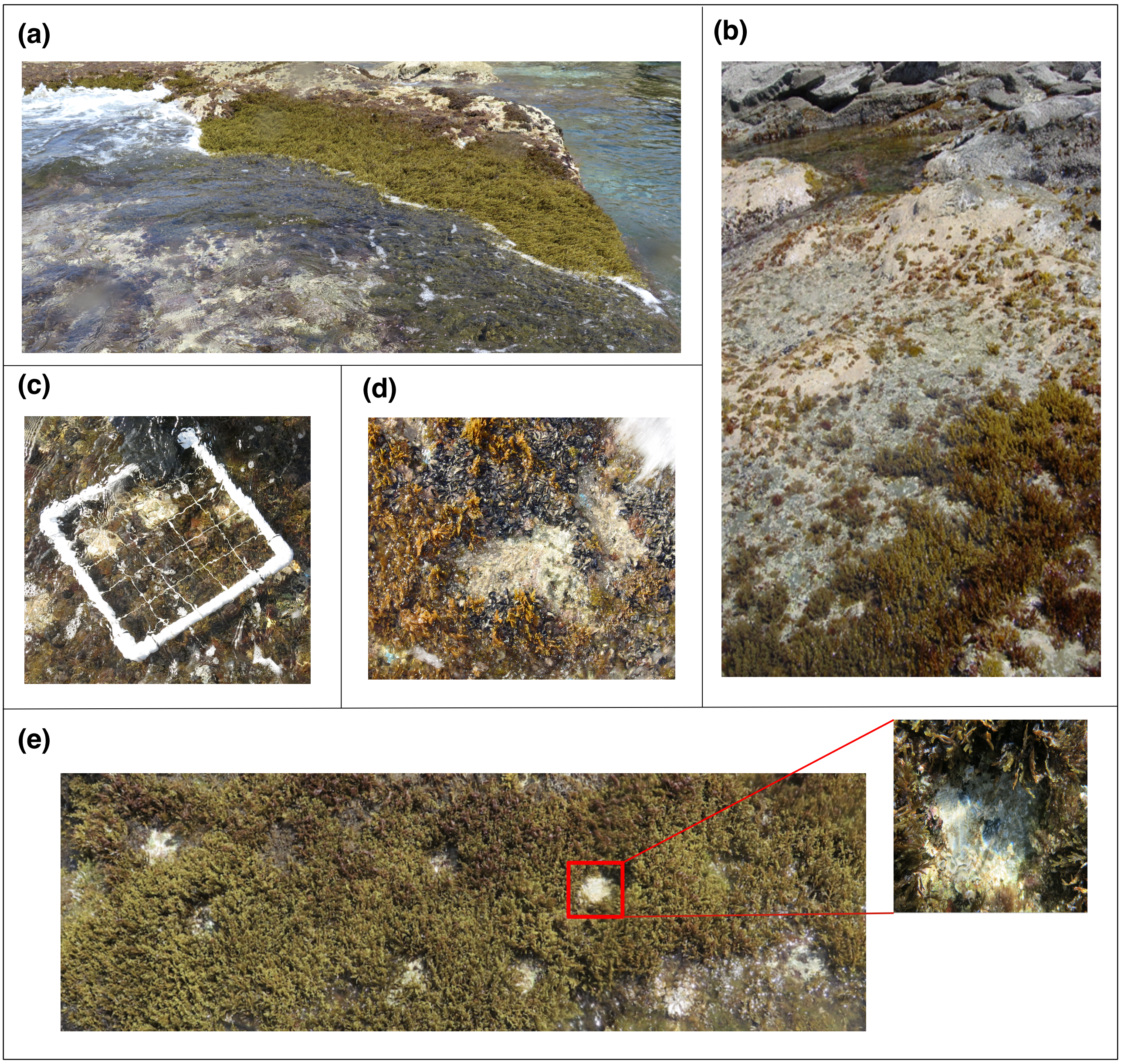


**Figure A1 Study system**. Images of the study system: (a) a stretch of coast fully covered by *Cystoseira compressa*. (b) naturally fragmented area. (c) plastic frame used for locating experimental gaps and used in high-resolution sampling. (d) a close-up of clumps of mussels (*Mytilus galloprovincialis*) within a patch of *C. compressa*. (e) a view of experimental clearings within a HOM-L transect. The inset shows a close-up of an experimental gap.

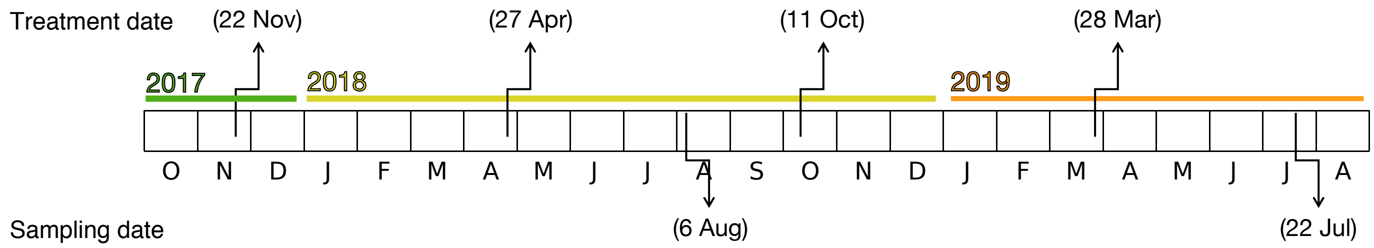


**Figure A2 Timeline of the experiment.** Transects were disturbed four times over the course of experiment (from November 2017 to July 2019). All transects were sampled in August 2018 and July 2019, about four months after the second and the fourth treatment.

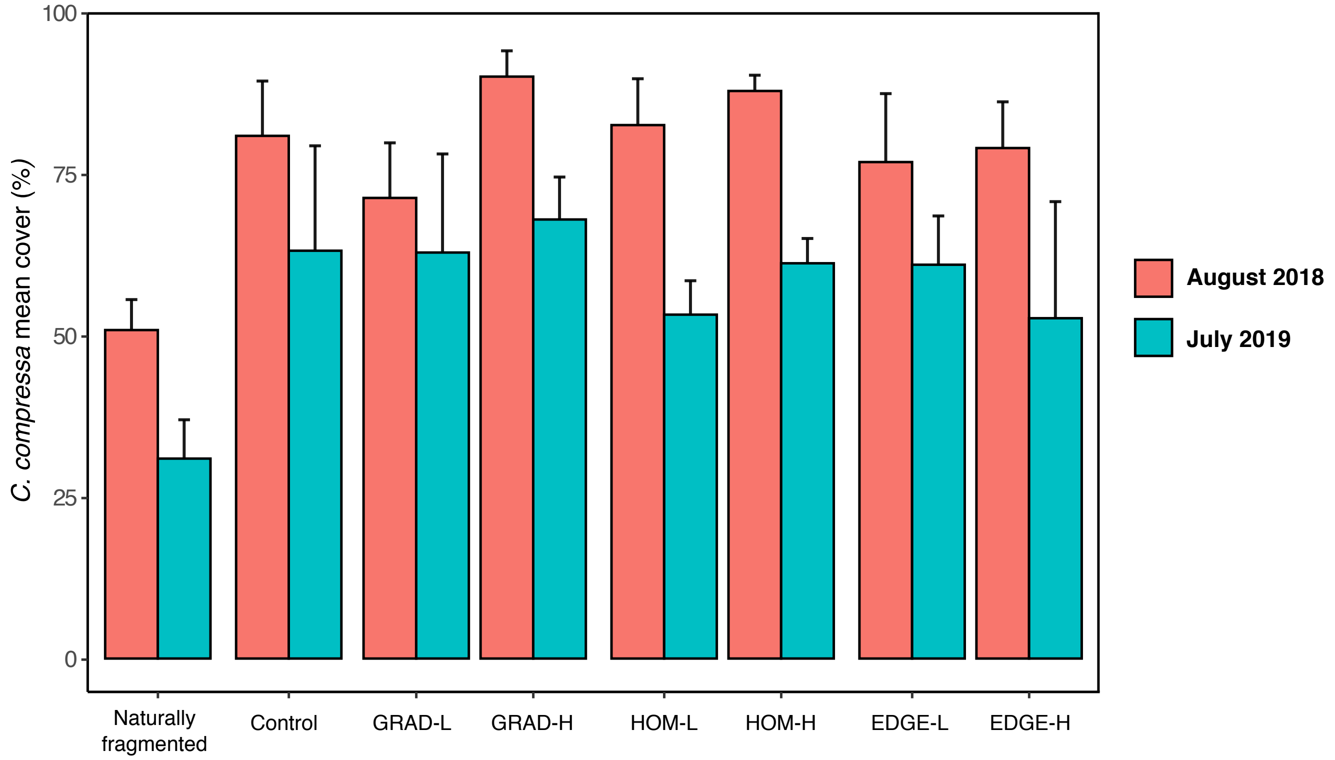


**Figure A3 Percentage cover of *C. compressa* (mean +SE) under different experimental conditions at different times.**

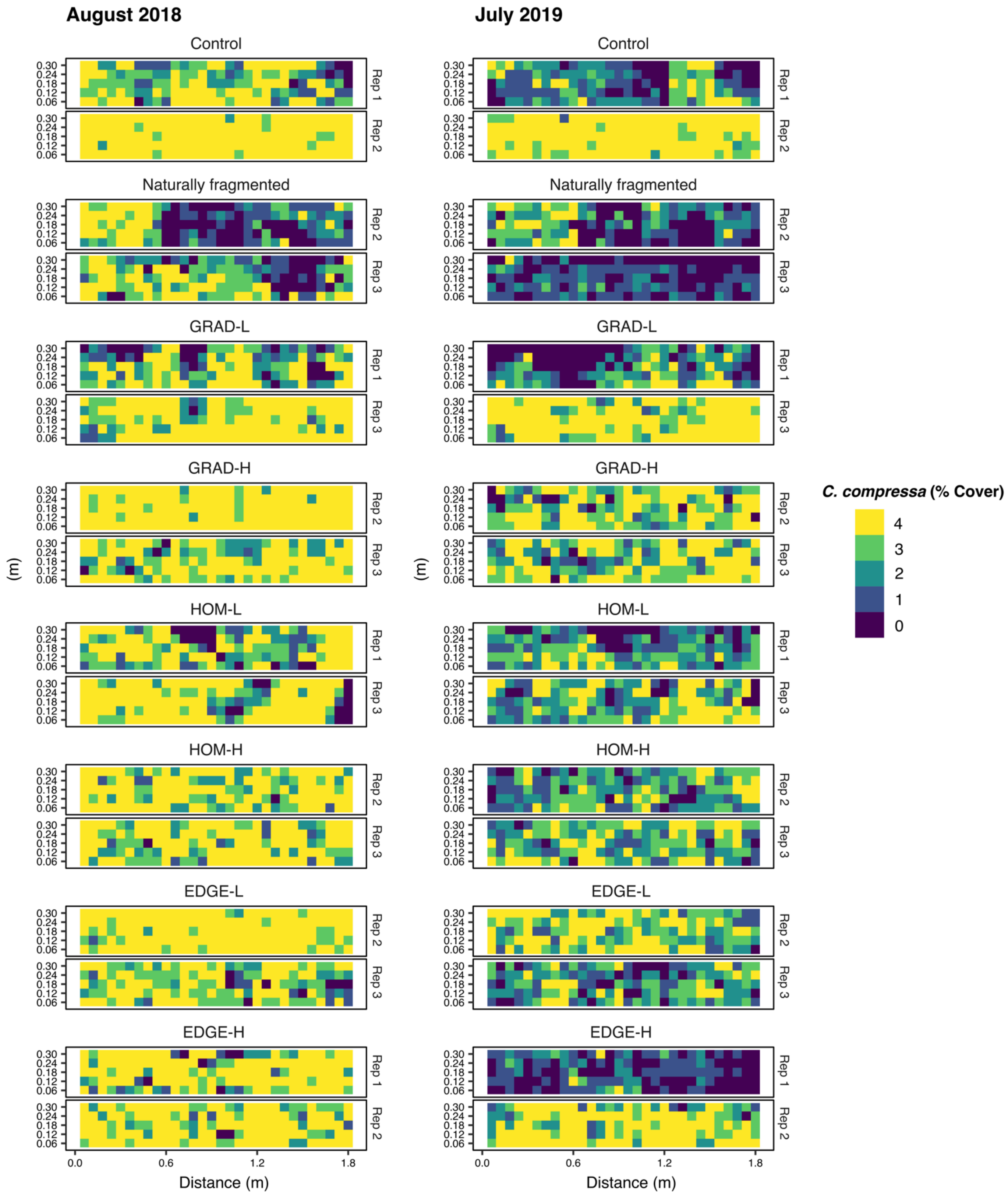


**Figure A4 Heat-maps of *C. compressa* cover in August 2018 and July 2019.** Two replicates for each treatment are shown here; one replicate is presented in Fig. 4 of main text.

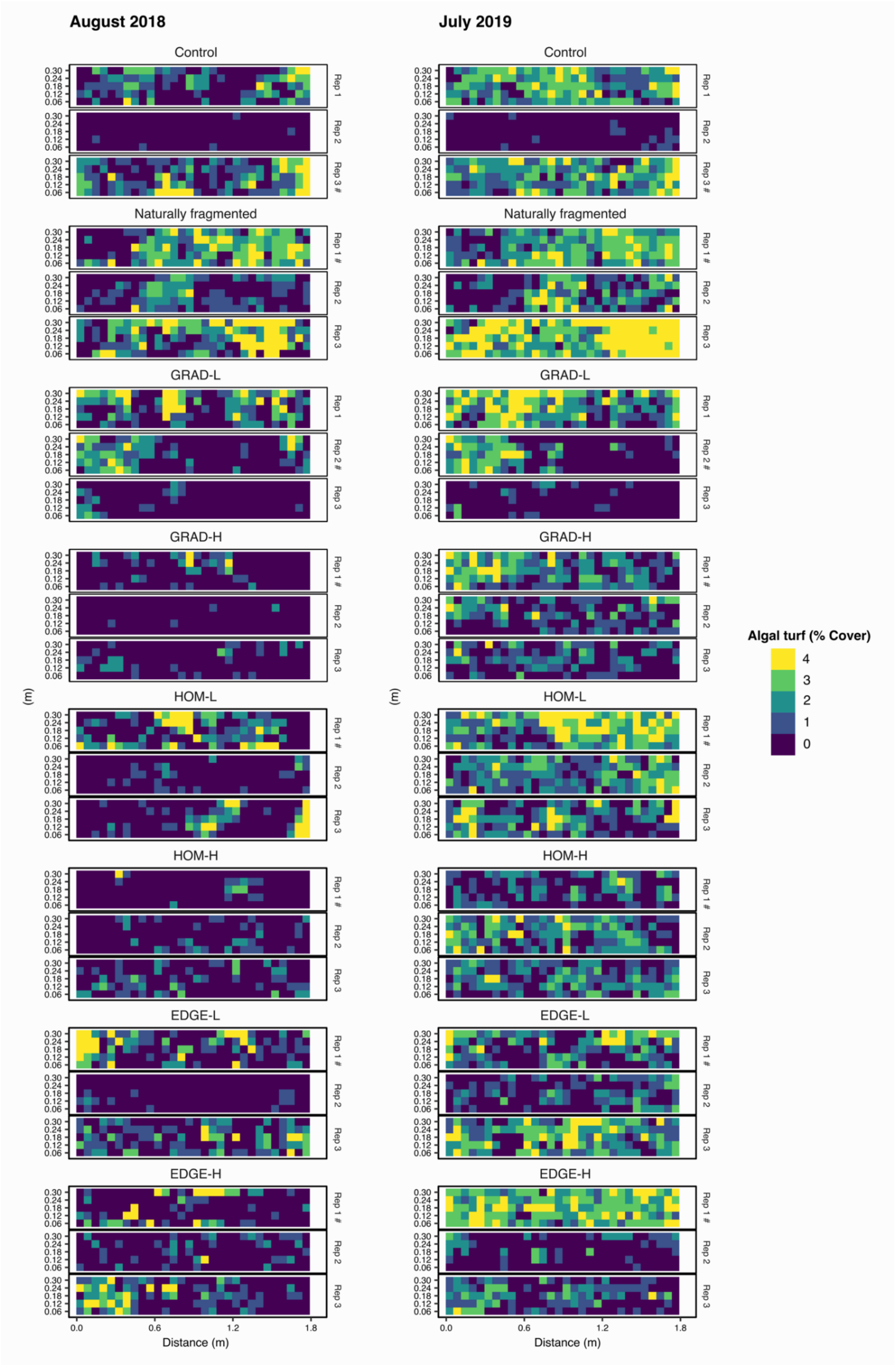


**Figure A5 Heat-maps of algal turfs cover in August 2018 and July 2019.** Experimental transects presented in Figure 4 are marked with a hashtag.

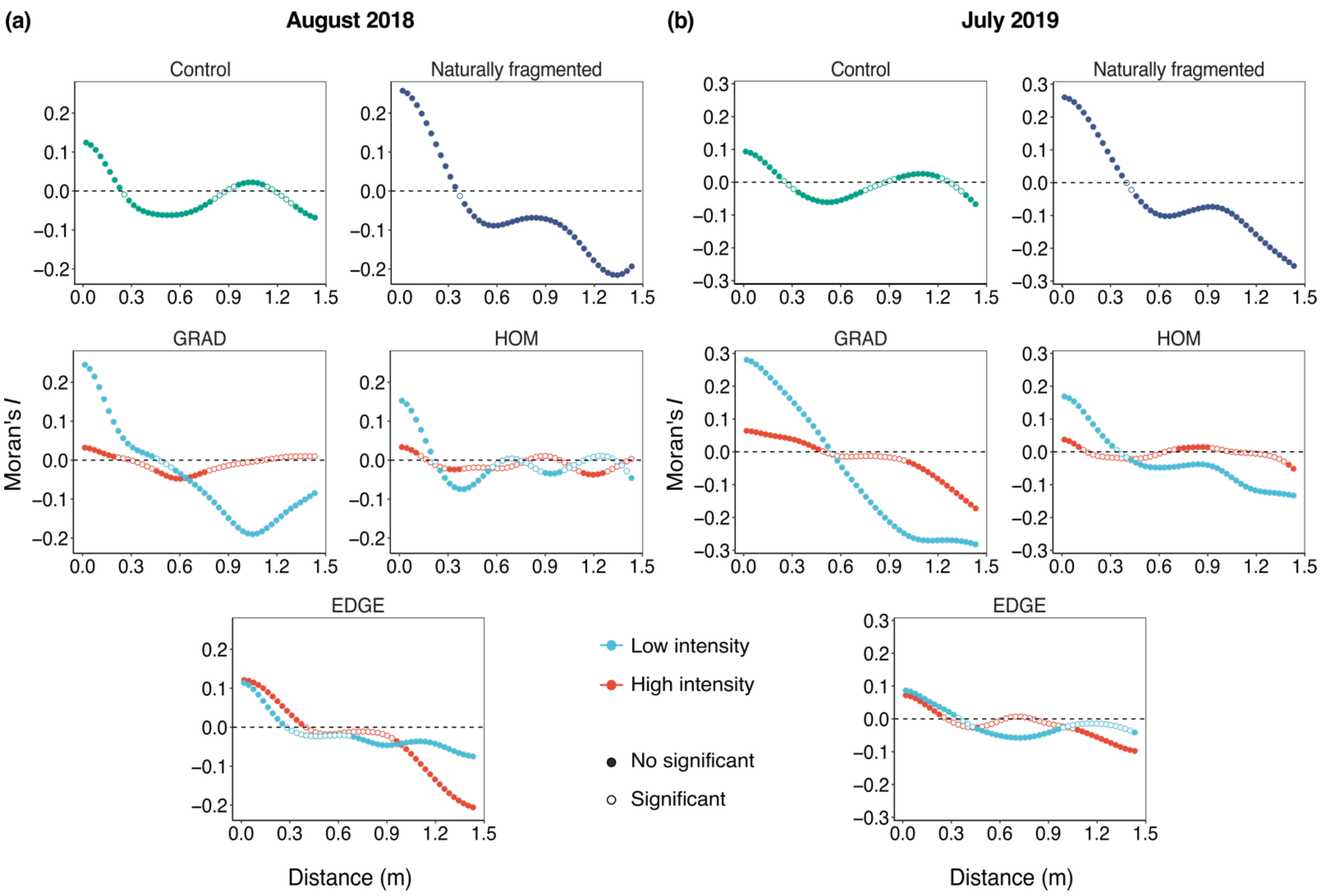


**Figure A6 Moran's *I* correlograms of *algal turfs* cover.** Spatial autocorrelation (Moran's *I*) as a function of distance in (a) August 2018 and (b) July 2019. Significant values of Moran's *I* (p 0.05) are denoted by filled symbols. Moran's *I* significance at each distance was tested over 1,000 permutations. Note that in order to improve readability, the y-axis limits are rescaled according to observed values of Moran's *I*.

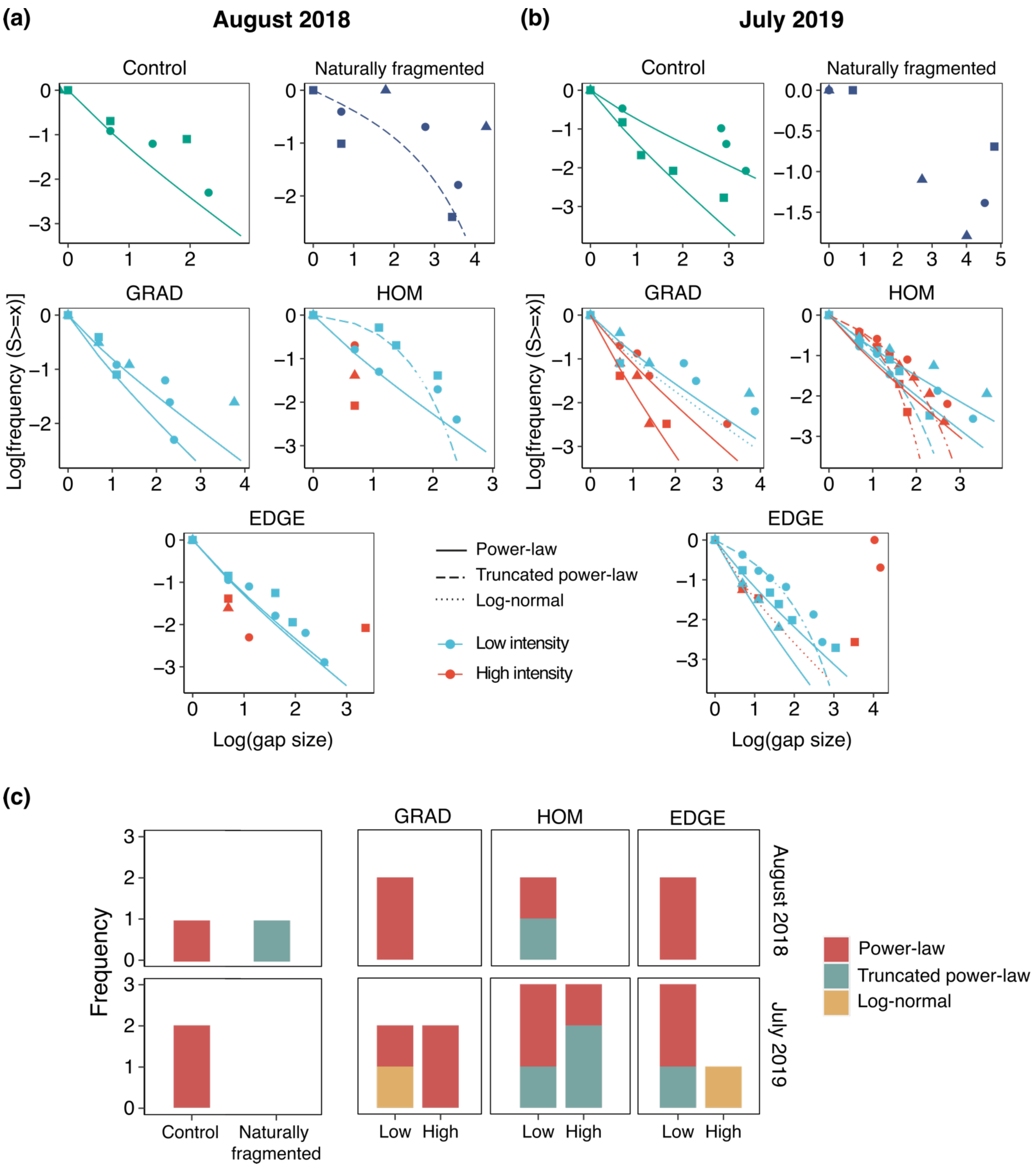


**Figure A7 Frequency distributions of gap sizes in experimental transects.** (a,b) Plots of log-inverse cumulative distribution of gap sizes - log-transformed frequency of gaps larger than a certain size (*s*) - as a function of the log of gap size. Lines denote the three alternative distributions: power-law (continuous), truncated power-law (dashed) and log-normal (dotted). Symbol shapes denote replicate transects (n=3). Transects with less than four distinct gap sizes were excluded from the analysis. (c) Absolute frequency of selected distributions (among those tested) in different experimental conditions in August 2018 and July 2019. Colors denote the three alternative distributions: power-law (green), truncated power-law (yellow) and log-normal (red).

**Table A2 Results of model fit on patch-size distribution.** Three alternative distributions (power-law, truncated power-law and log-normal) were fitted to patch-size data of algal turfs. Mean and standard deviation refer to the log-normal distribution. The optimal distribution was selected on the basis of the lowest Akaike Information Criterion (AIC) (blue shadowing). PL: Power-law distribution; TPL: Truncated power-law distribution; LNORM: Lognormal distribution.

| Treatment | Transect | Function | AIC | Mean | SD | Scaling exponent (λ) | Exponential decay rate (β) |
|---|---|---|---|---|---|---|---|
| **August 2018** | | | | | | | |
| Naturally fragmented | 1 | PL | 50.59 | - | - | 3.55 | - |
| | | TPL | 49.61 | - | - | 2.38 | 5.81 |
| | | LNORM | 50.93 | 0.73 | 1.72 | - | - |
| | 3 | PL | 47.05 | - | - | 3.46 | - |
| | | TPL | 47.24 | - | - | 2.23 | 6.21 |
| | | LNORM | 47.42 | 0.74 | 1.51 | - | - |
| Control | 1 | PL | 35.75 | - | - | 2.92 | - |
| | | TPL | 36.10 | - | - | 2.33 | 4.26 |
| | | LNORM | 36.29 | 0.18 | 1.01 | - | - |
| | 3 | PL | 38.64 | - | - | 3.23 | - |
| | | TPL | 39.39 | - | - | 2.17 | 5.67 |
| | | LNORM | 39.87 | -0.28 | 1.73 | - | - |
| GRAD-L | 1 | PL | 46.74 | - | - | 3.21 | - |
| | | TPL | 45.53 | - | - | 2.54 | 4.52 |
| | | LNORM | 46.23 | 0.61 | 1.10 | - | - |
| | 2 | PL | 27.18 | - | - | 2.98 | - |
| | | TPL | 28.21 | - | - | 2.21 | 4.75 |
| | | LNORM | 28.56 | -0.27 | 1.36 | - | - |
| HOM-L | 1 | PL | 39.40 | - | - | 3.02 | - |
| | | TPL | 40.01 | - | - | 2.19 | 4.90 |
| | | LNORM | 40.27 | 0.02 | 1.25 | - | - |
| | 3 | PL | 23.22 | - | - | 3.39 | - |
| | | TPL | 21.78 | - | - | 4.52 | 3.75 |
| | | LNORM | 22.23 | 1.13 | 0.77 | - | - |
| EDGE-L | 1 | PL | 35.21 | - | - | 2.66 | - |
| | | TPL | 37.18 | - | - | 1.72 | 7.63 |
| | | LNORM | 31.98 | -1256.64 | 33.56 | - | - |
| | 3 | PL | 30.05 | - | - | 2.65 | - |
| | | TPL | 31.05 | - | - | 2.19 | 3.97 |
| | | LNORM | 31.29 | -0.29 | 1.02 | - | - |
| EDGE-H | 1 | PL | 27.97 | - | - | 2.77 | - |
| | | TPL | 28.70 | - | - | 2.37 | 3.89 |
| | | LNORM | 28.82 | 0.08 | 0.91 | - | - |
| | 3 | PL | 31.27 | - | - | 3.00 | - |

| | | | | | | | |
|---|---|---|---|---|---|---|---|
| | | TPL | 31.59 | - | - | 2.42 | 4.27 |
| | | LNORM | 32.02 | 0.23 | 1.08 | - | - |
| **July 2019** | | | | | | | |
| Naturally fragmented | 1 | PL | 59.67 | - | - | 3.12 | - |
| | | TPL | 61.04 | - | - | 1.97 | 6.85 |
| | | LNORM | 61.32 | -5.00 | 3.03 | - | - |
| | 2 | PL | 35.20 | - | - | 2.81 | - |
| | | TPL | 37.20 | - | - | 1.76 | 16.10 |
| | | LNORM | 36.87 | -70.27 | 8.56 | - | - |
| | 3 | PL | 47.24 | - | - | 3.33 | - |
| | | TPL | 49.05 | - | - | 1.97 | 9.21 |
| | | LNORM | 39.63 | -2286.23 | 61.05 | - | - |
| Control | 1 | PL | 62.23 | - | - | 3.25 | - |
| | | TPL | 62.48 | - | - | 2.13 | 5.95 |
| | | LNORM | 62.77 | 0.10 | 1.54 | - | - |
| | 3 | PL | 70.96 | - | - | 3.24 | - |
| | | TPL | 65.05 | - | - | 3.18 | 3.90 |
| | | LNORM | 64.63 | 0.88 | 0.78 | - | - |
| GRAD-L | 1 | PL | 41.28 | - | - | 3.31 | - |
| | | TPL | 42.92 | - | - | 2.00 | 7.91 |
| | | LNORM | 43.09 | -1.65 | 2.44 | - | - |
| GRAD-H | 1 | PL | 35.49 | - | - | 2.91 | - |
| | | TPL | 37.38 | - | - | 1.85 | 7.41 |
| | | LNORM | 37.45 | -5.94 | 2.89 | - | - |
| HOM-L | 1 | PL | 45.04 | - | - | 3.41 | - |
| | | TPL | 46.42 | - | - | 2.05 | 7.65 |
| | | LNORM | 46.29 | 0.18 | 1.81 | - | - |
| | 2 | PL | 48.60 | - | - | 3.06 | - |
| | | TPL | 48.84 | - | - | 2.20 | 5.00 |
| | | LNORM | 49.20 | 0.04 | 1.29 | - | - |
| | 3 | PL | 41.41 | - | - | 2.89 | - |
| | | TPL | 42.62 | - | - | 2.01 | 5.29 |
| | | LNORM | 42.90 | -1.01 | 1.58 | - | - |
| HOM-H | 2 | PL | 46.33 | - | - | 2.79 | - |
| | | TPL | 47.57 | - | - | 1.96 | 5.17 |
| | | LNORM | 47.88 | -2.04 | 1.80 | - | - |
| EDGE-L | 1 | PL | 44.92 | - | - | 2.98 | - |
| | | TPL | 45.76 | - | - | 2.09 | 5.17 |
| | | LNORM | 46.26 | -0.93 | 1.65 | - | - |
| | 3 | PL | 54.98 | - | - | 3.39 | - |
| | | TPL | 54.38 | - | - | 2.29 | 5.65 |
| | | LNORM | 54.99 | 0.64 | 1.44 | - | - |
| EDGE-H | 1 | PL | 48.89 | - | - | 3.84 | - |
| | | TPL | 47.58 | - | - | 2.43 | 6.63 |

| | | | | | | | |
|---|---|---|---|---|---|---|---|
| | | LNORM | 48.98 | 1.50 | 1.90 | - | - |
| | 3 | PL | 26.16 | - | - | 2.61 | - |
| | | TPL | 27.68 | - | - | 1.96 | 4.52 |
| | | LNORM | 27.87 | -1.31 | 1.39 | - | - |

**Table A2 Results of model fit on gap-size distribution.** Three alternative distributions (power-law, truncated power-law and log-normal) were fitted to the gap-size distribution of *Cystoseira compressa*. Mean and standard deviation refer to the log-normal distribution. The optimal distribution was selected based on the lowest Akaike Information Criterion (AIC) (blue shadowing). PL: Power-law distribution; TPL: Truncated power-law distribution; LNORM: Lognormal distribution.

| Treatment | Transect | Function | AIC | Mean | SD | Scaling exponent (λ) | Exponential decay rate (β) |
|---|---|---|---|---|---|---|---|
| **August 2018** | | | | | | | |
| Naturally fragmented | 1 | PL | 42.96 | - | - | 3.68 | - |
| | | TPL | 42.88 | - | - | 2.30 | 6.63 |
| | | LNORM | 43.84 | 0.74 | 2.05 | - | - |
| Control | 1 | PL | 35.03 | - | - | 2.90 | - |
| | | TPL | 36.35 | - | - | 2.01 | 5.29 |
| | | LNORM | 36.59 | -1.08 | 1.61 | - | - |
| GRAD-L | 1 | PL | 43.44 | - | - | 3.13 | - |
| | | TPL | 44.53 | - | - | 2.06 | 5.95 |
| | | LNORM | 44.97 | -2.48 | 2.38 | - | - |
| | 2 | PL | 28.96 | - | - | 3.41 | - |
| | | TPL | 30.63 | - | - | 2.04 | 7.94 |
| | | LNORM | 30.83 | -2.79 | 3.01 | - | - |
| HOM-L | 1 | PL | 40.75 | - | - | 2.96 | - |
| | | TPL | 42.00 | - | - | 2.01 | 5.56 |
| | | LNORM | 42.27 | -0.96 | 1.65 | - | - |
| | 3 | PL | 23.22 | - | - | 3.39 | - |
| | | TPL | 21.78 | - | - | 4.52 | 3.75 |
| | | LNORM | 22.23 | 1.13 | 0.77 | - | - |
| EDGE-L | 1 | PL | 61.36 | - | - | 2.90 | - |
| | | TPL | 62.52 | - | - | 1.95 | 5.73 |
| | | LNORM | 62.81 | -1.77 | 1.85 | - | - |
| | 3 | PL | 25.95 | - | - | 2.93 | - |
| | | TPL | 27.16 | - | - | 2.15 | 4.79 |
| | | LNORM | 27.46 | -0.48 | 1.40 | - | - |
| **July 2019** | | | | | | | |
| Control | 1 | PL | 48.88 | - | - | 3.50 | - |
| | | TPL | 49.42 | - | - | 2.17 | 6.80 |
| | | LNORM | 50.19 | -0.49 | 2.34 | - | - |
| | 3 | PL | 52.12 | - | - | 2.85 | - |
| | | TPL | 53.79 | - | - | 1.86 | 6.45 |
| | | LNORM | 53.89 | -1.68 | 1.77 | - | - |
| GRAD-L | 1 | PL | 44.04 | - | - | 3.25 | - |
| | | TPL | 45.82 | - | - | 1.96 | 8.50 |

| | | | | | | | |
|---|---|---|---|---|---|---|---|
| | | LNORM | 37.01 | -2110.66 | 56.36 | - | - |
| | 2 | PL | 32.73 | - | - | 3.35 | - |
| | | TPL | 34.38 | - | - | 2.02 | 7.78 |
| | | LNORM | 34.57 | -1.38 | 2.43 | - | - |
| GRAD-H | 1 | PL | 49.13 | - | - | 3.07 | - |
| | | TPL | 50.55 | - | - | 1.97 | 6.52 |
| | | LNORM | 50.58 | -0.85 | 1.75 | - | - |
| | 2 | PL | 30.29 | - | - | 2.58 | - |
| | | TPL | 30.83 | - | - | 2.50 | 3.38 |
| | | LNORM | 31.22 | -0.12 | 0.86 | - | - |
| HOM-L | 1 | PL | 54.65 | - | - | 3.10 | - |
| | | TPL | 56.15 | - | - | 1.95 | 6.98 |
| | | LNORM | 56.35 | -2.82 | 2.46 | - | - |
| | 2 | PL | 39.45 | - | - | 3.40 | - |
| | | TPL | 40.73 | - | - | 2.08 | 7.16 |
| | | LNORM | 41.05 | -1.22 | 2.45 | - | - |
| | 3 | PL | 48.45 | - | - | 3.06 | - |
| | | TPL | 48.11 | - | - | 2.33 | 4.59 |
| | | LNORM | 48.74 | 0.18 | 1.19 | - | - |
| HOM-H | 1 | PL | 46.16 | - | - | 3.31 | - |
| | | TPL | 45.08 | - | - | 2.49 | 4.84 |
| | | LNORM | 45.40 | 0.79 | 1.11 | - | - |
| | 2 | PL | 55.21 | - | - | 3.04 | - |
| | | TPL | 56.40 | - | - | 1.99 | 6.13 |
| | | LNORM | 56.70 | -4.25 | 2.71 | - | - |
| | 3 | PL | 47.08 | - | - | 3.12 | - |
| | | TPL | 43.21 | - | - | 3.92 | 3.41 |
| | | LNORM | 44.08 | 0.72 | 0.75 | - | - |
| EDGE-L | 1 | PL | 66.05 | - | - | 3.31 | - |
| | | TPL | 63.77 | - | - | 2.46 | 4.92 |
| | | LNORM | 64.20 | 0.78 | 1.13 | - | - |
| | 2 | PL | 24.11 | - | - | 2.62 | - |
| | | TPL | 25.44 | - | - | 2.12 | 4.07 |
| | | LNORM | 25.63 | -0.50 | 1.09 | - | - |
| | 3 | PL | 57.03 | - | - | 3.00 | - |
| | | TPL | 58.38 | - | - | 1.95 | 6.30 |
| | | LNORM | 58.57 | -1.43 | 1.87 | - | - |
| EDGE-H | 3 | PL | 40.52 | - | - | 2.79 | - |
| | | TPL | 42.51 | - | - | 1.76 | 11.42 |
| | | LNORM | 35.17 | -1419.51 | 37.93 | - | - |
